## Supplementary material for "Community composition of microbial microcosms follows simple assembly rules at evolutionary timescales"

### Supplementary Information

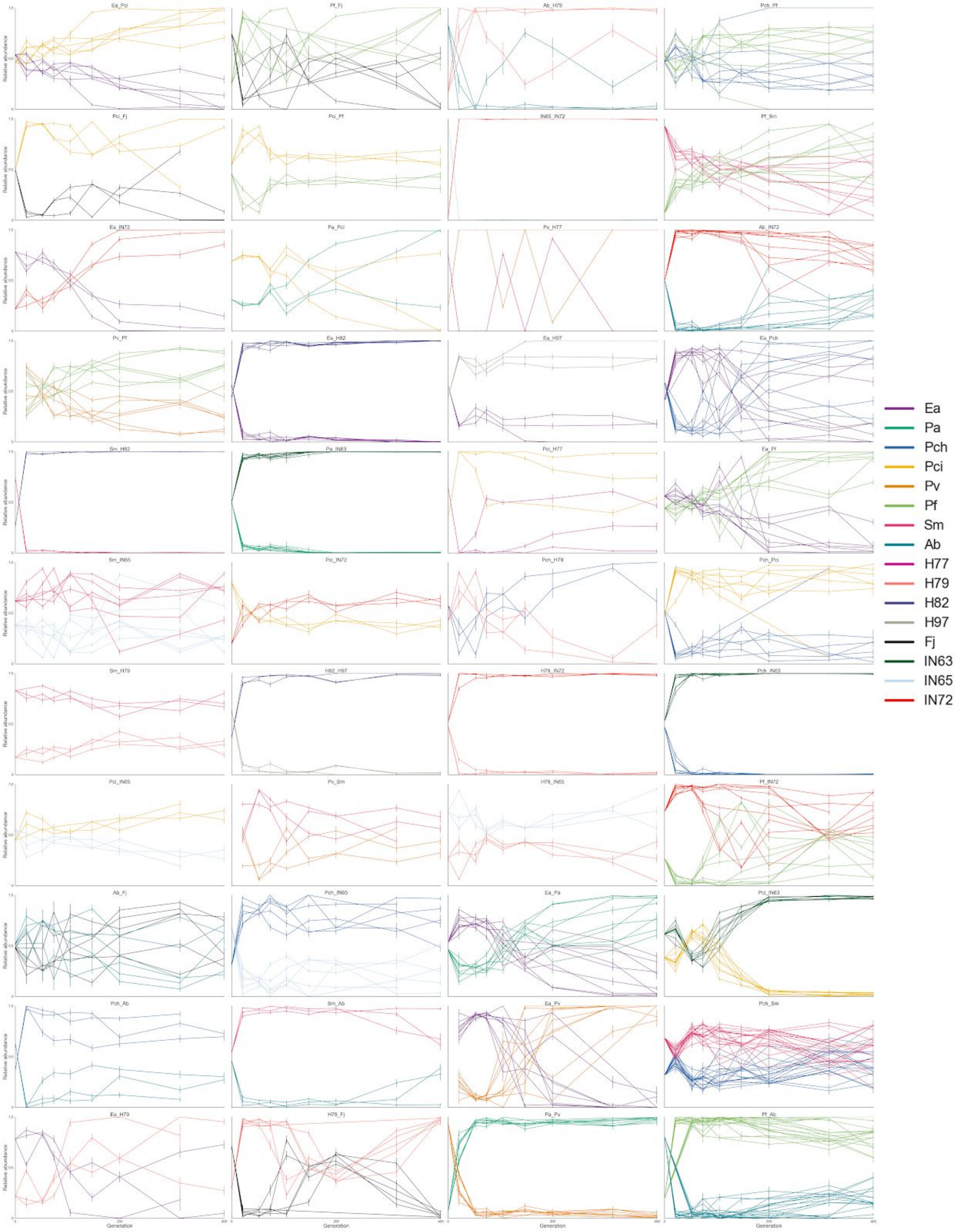


***Figure S1: The community composition of all pairs throughout the experiment.*** *Colors denote the different species, and lines are replicates. The community composition was assayed by plating on agar plates and counting colonies, which are distinct for each species. Error bars indicate the SD derived from binomial distribution.*

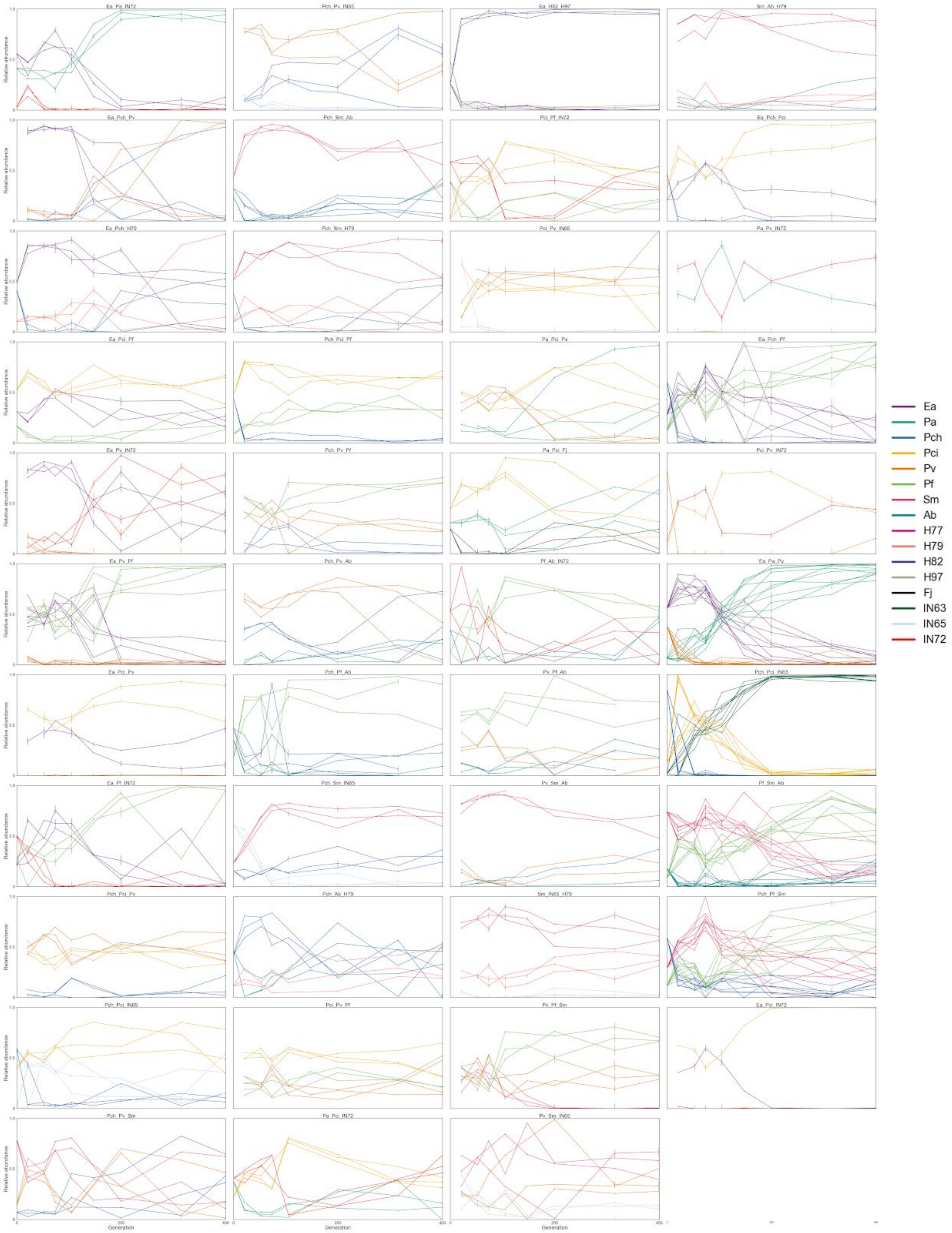


***Figure S2: The community composition of all trios throughout the experiment.*** *Colors denote the different species, and lines are replicates. The community composition was assayed by plating on agar plates and counting colonies, which are distinct for each species.*

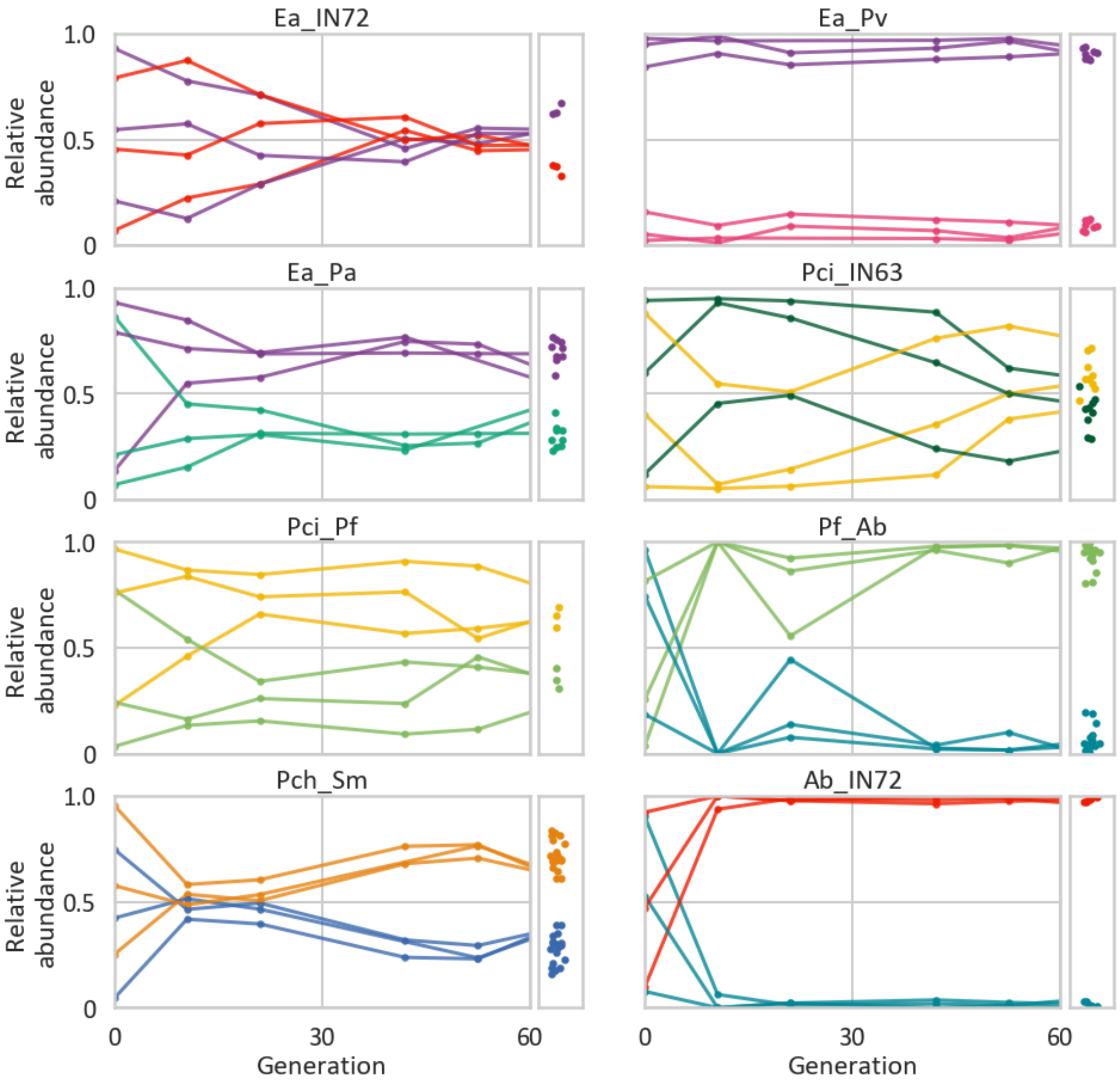


***Figure S3: Pairs converge to the same composition from multiple initial fractions.*** *Line plots indicate the composition of pairs that started a ~60 generation competition experiment from three different initial conditions. Species were normalized to three initial fractions (0.1, 0.5, 0.9) by OD, thus a difference in the actual initial fractions as measured by CFU is to be expected. Colors denote different species. Narrow scatter plots near each line plot, indicate the fractions of the same pairs at the separate ~400 generation evolutionary experiment at generation ~70.*

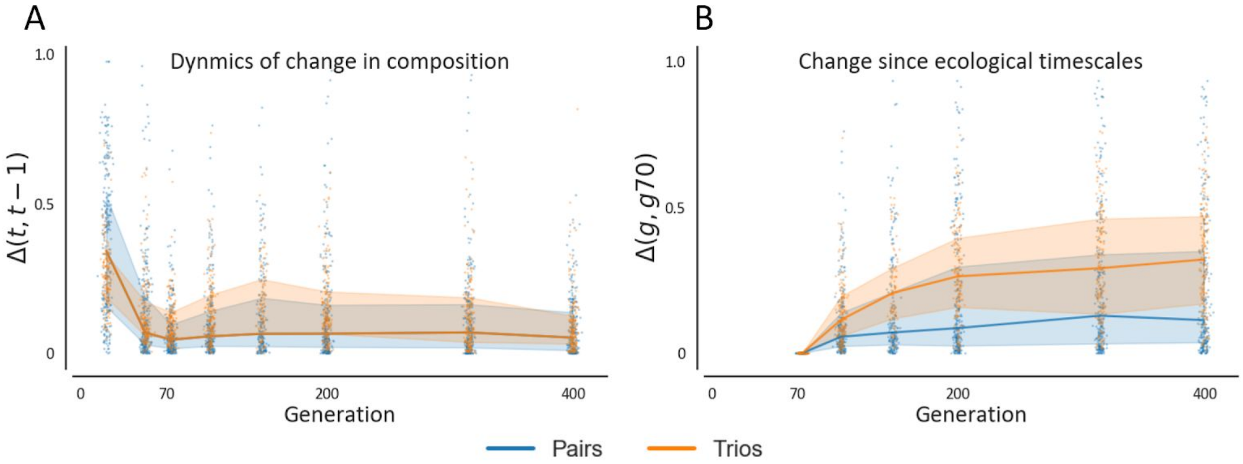


***Figure S4: Dynamics of change in composition of pairs and trios separately.*** *(A) Change in community composition across all pairs and trios quantified as the Euclidean distance between the composition of each replicate at two subsequent time points normalized to the maximal distance between two communities composed of n species (*$\sqrt{n},$*), denoted as* $\Delta(t,\ t-1)$*. (A) Change in community composition at evolutionary timescales measured as the Euclidean distance between the composition of each replicate in each time point and its composition at generation ~70 (*$\Delta(g,\ g_{70})$*). Generation ~70 is used here as the starting point of the evolutionary timescale since changes in most communities are less rapid after these timescales. For both B and C, lines denote the median, and shaded areas denote the interquartile range across all communities.*

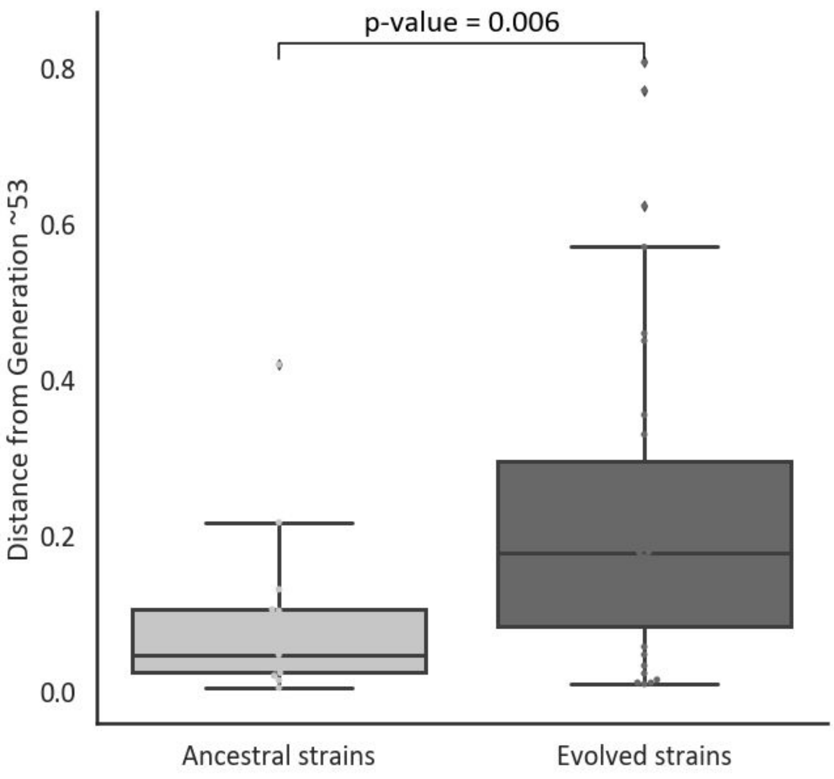


***Figure S5. Pairs of evolved strains reach different compositions than those reached by their ancestors.*** *Data indicate the Euclidean distance of the composition reached by ancestral strains, and reisolated evolved strains in a short ~53 generation experiment, to the composition that was reached at the evolutionary experiment at generation ~53. Dots of ancestral strains indicate the mean of three technical replicates. Dots of evolved strains are averaged across two technical replicates, and 1-3 evolutionary replicates (communities that evolved in different wells). Boxes*

*indicate the interquartile range and whiskers are expanded to include values no further than 1.5X interquartile range. P-value indicates the p-value obtained by a Mann-Whitney U-test.*

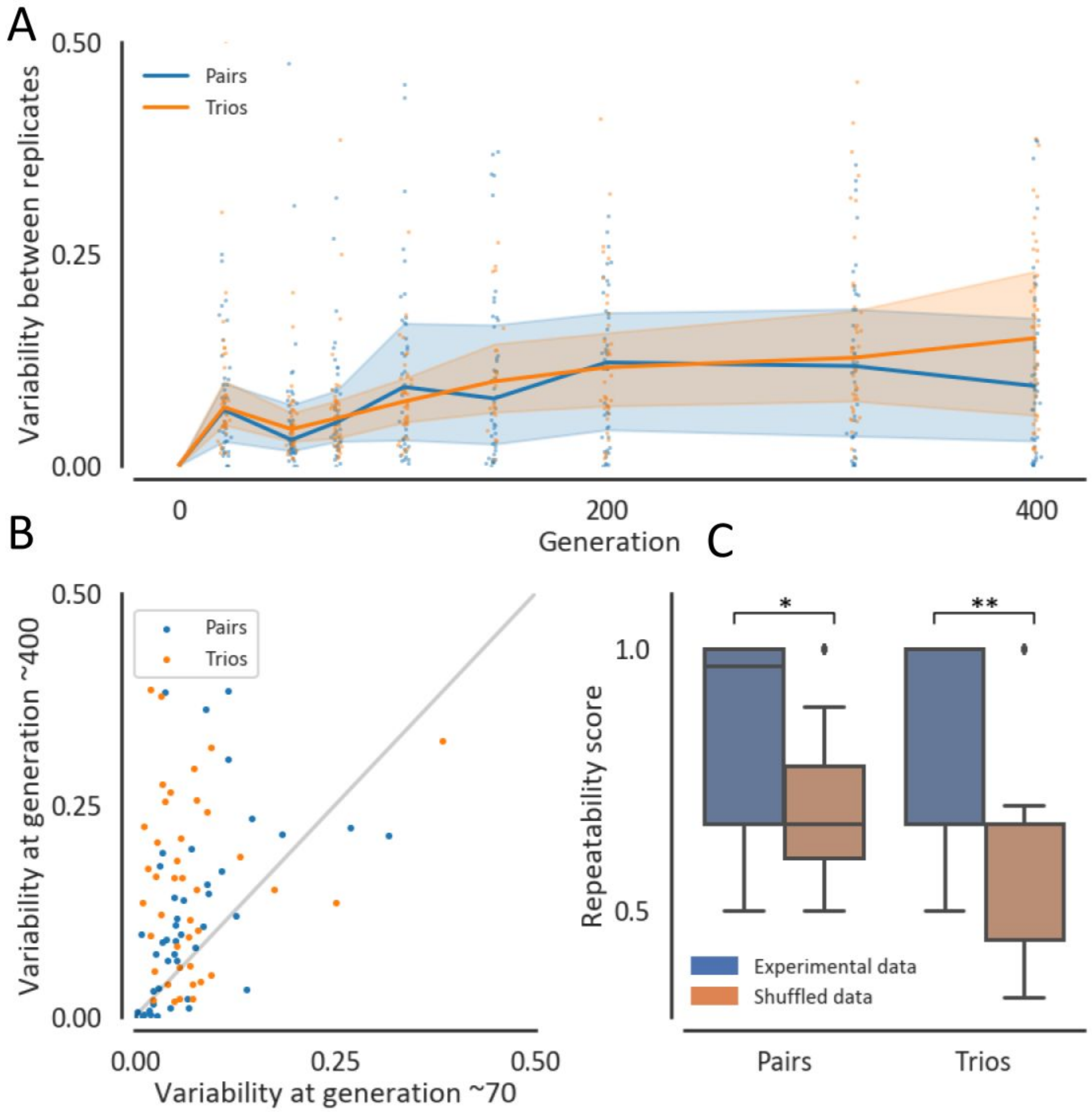


***Figure S6: Repeatability of pairs and trios.*** *(A) Variability in community composition between replicate communities is quantified by the mean Euclidean distance of each replicate from the medoid replicate normalized to the maximal distance between two communities composed of n species ($\sqrt{n}$, ). Blue and orange line and shaded-area are the median and interquartile range across all pairs and trios respectively. Dots denote the variability of specific pairs and trios across replicates (B) Variability at generation ~400 against ~70, measured as the mean distance from. Each dot represents a pair (blue) or a trio (orange). (C) Distribution of repeatability scores of the experimental data and a shuffled null model. The repeatability score is the frequency of replicates in which the same species increased its abundance by the biggest factor between generation ~70 and ~400. The brown box represents the distribution of a fully shuffled model, where species increase values are pooled and are randomly assigned to any species in any community in the dataset. Boxes indicate the interquartile range and whiskers are expanded to include values no further than 1.5X interquartile range. * p-value=0.05, **p-value<0.005*

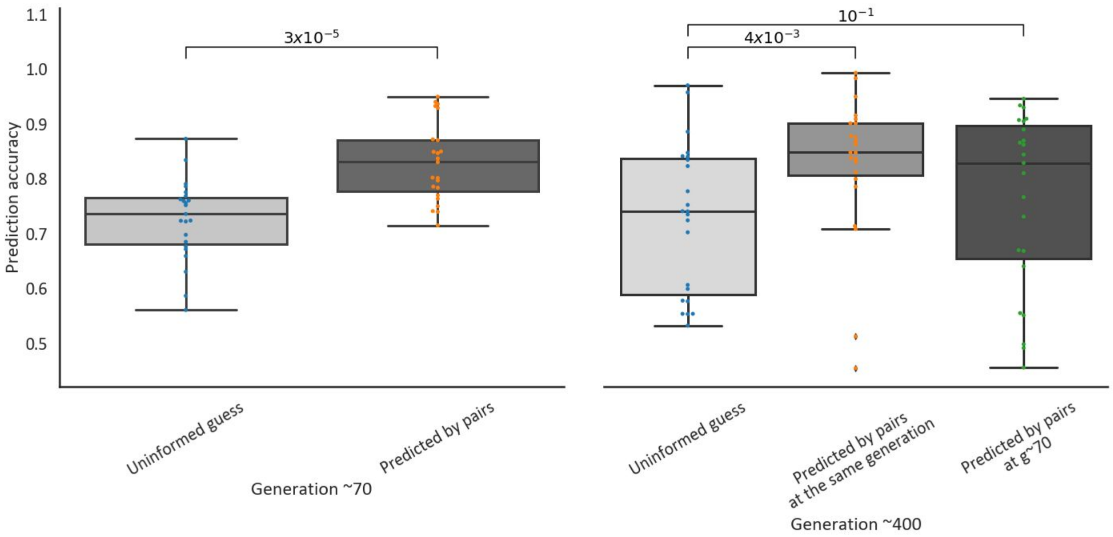


***Figure S7: The composition of pairs predicts trios.*** *The distribution of prediction accuracies made by using the composition of pairs. The prediction accuracy is measured as* $1 - \frac{\Delta(Prediction,\, Observation)}{\sqrt{n}}$*, where* $\Delta(Prediction,\, Observation)$ *is the Euclidean distance between the prediction and the observation and* $\sqrt{n}$ *is the maximum distance between two communities with the same n species. Number above connectors indicate the p-values of a Mann-Whitney U test between the predictions and the null model. The null model is that all species have equal abundances in the trio. Boxes indicate the interquartile range and whiskers are expanded to include values no further then 1.5X interquartile range*

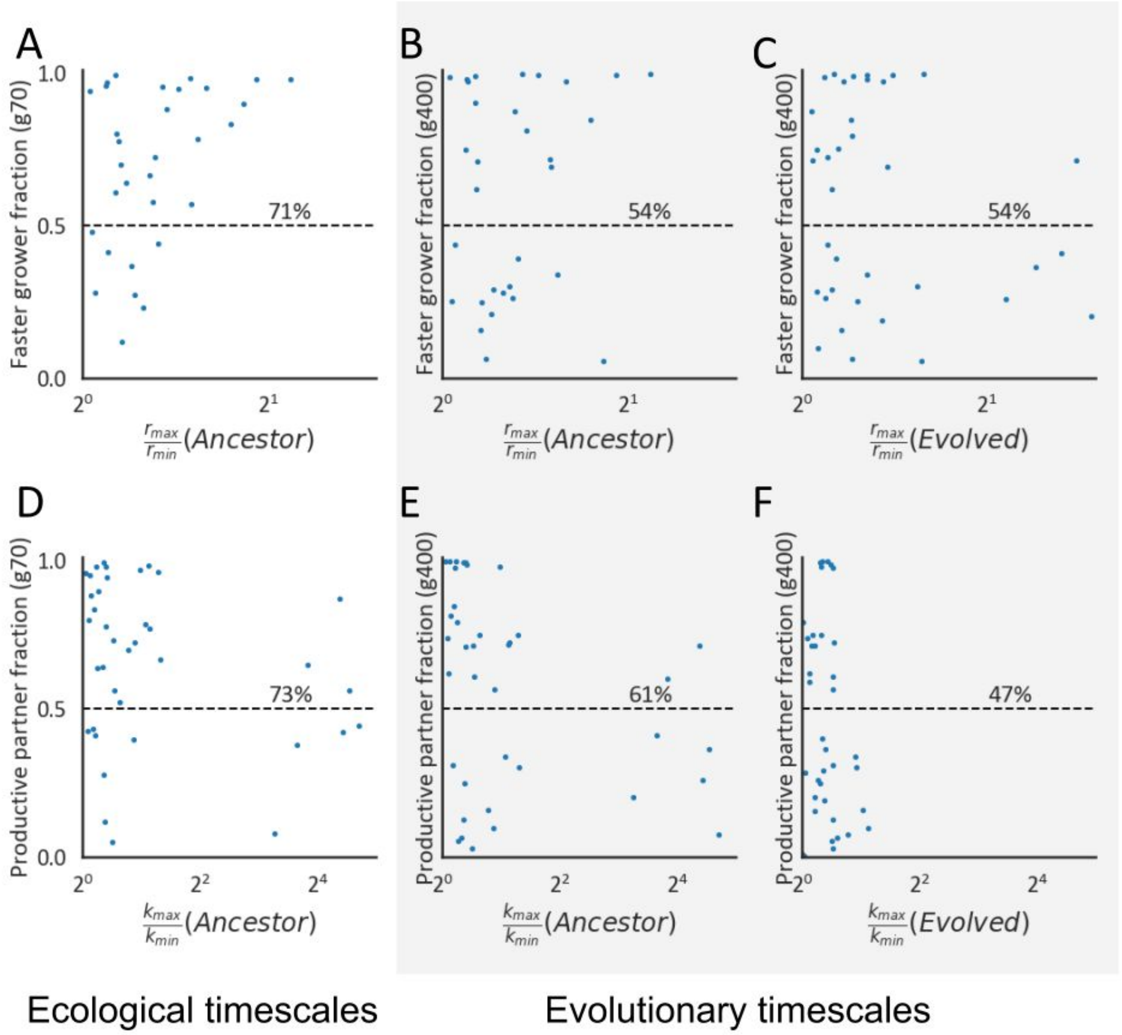


***Figure S8: Growth of strains that evolved in monoculture does not predict the composition of pairs.*** *(A-C) The mean fraction of the species with the higher growth rate (A and B – ancestor, C – evolved) at generation ~70 (A) and generation ~400 (B, C) vs the ratio of the higher growth rate and the lower growth rate. (D-E) The mean fraction of the species with the higher carrying capacity (D and E – ancestor, F – evolved) at generation ~70 (E) and generation ~400 (E, F) vs the ratio of the higher carrying capacity and the lower carrying capacity. Percentages indicate the percentage of pairs above the 0.5 line, which correspond to the prediction that the species with the higher growth rate/carrying capacity is more dominant in a pairwise competition.*

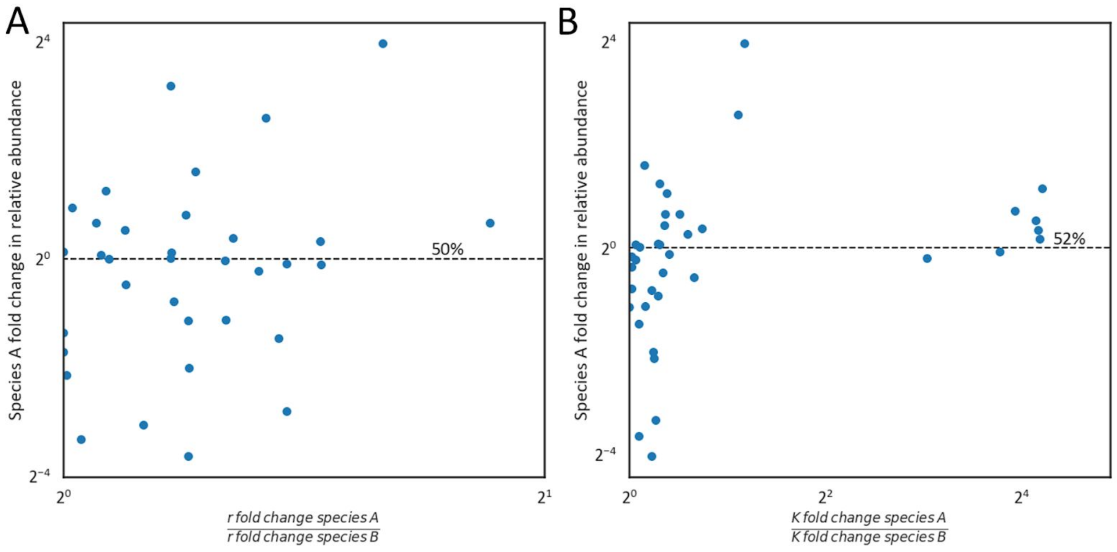


***Figure S9: The increase in growth parameters of monocultures does not predict the increase in relative abundance in pairs.*** *(A) the fold difference in increase in growth rate between species A and species B against A's fold change in relative abundance where A is always the species that increased in growth rate by the larger factor. (B) the fold difference in increase in carrying capacity between species A and species B against A's fold change in relative abundance where A is always the species that increased in carrying capacity by the larger factor. The increase in carrying capacity is measured as the mean OD of a monoculture in generation ~400 divided by the mean OD of a monoculture in generation ~70. For both A and B the percentages indicate the accuracy of the prediction that the species that increased its growth rate of carrying capacity by the larger factor is also the one that increased its relative abundance.*

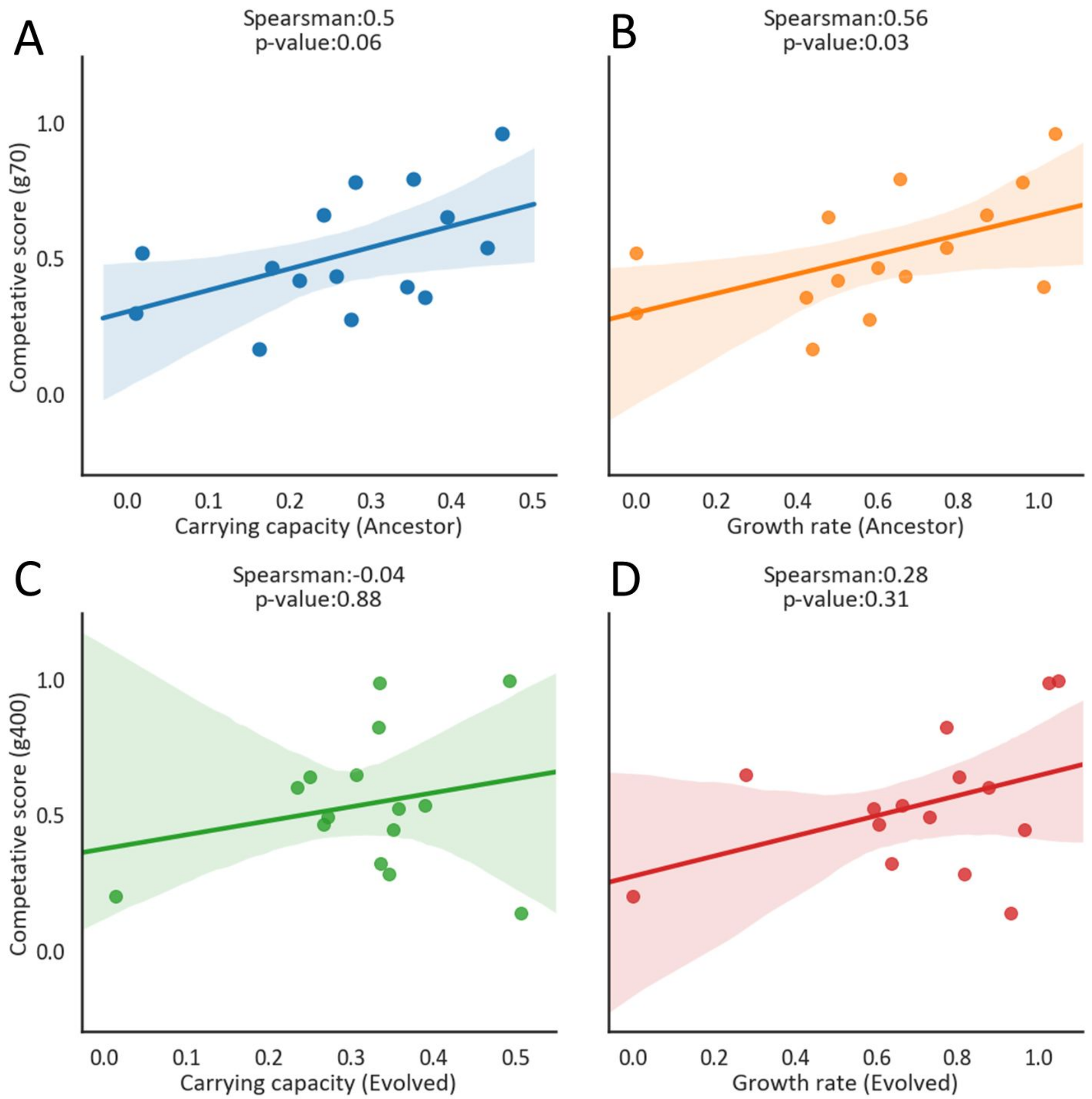


***Figure S10: Correlations between species' growth abilities and species' competitive scores.*** *(A) The ecological competitive score, as the mean fraction of a species at generation ~70, against its carrying capacity as its mean OD at generation ~70. (B) The ecological competitive score against the mean effective growth rate of the ancestor (Methods). (C) The evolutionary competitive score, as the mean fraction of a species at generation ~400, against its carrying capacity as its mean OD at generation ~400. (D) The evolutionary competitive score against the mean effective growth rate of the monoculturally evolved strains.*

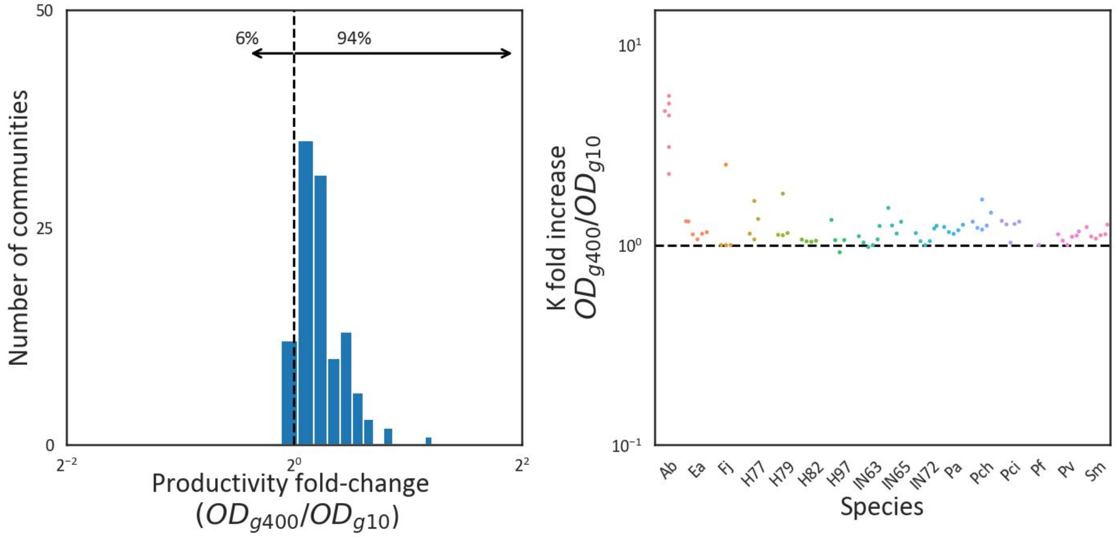


***Figure S11: Most communities increase their productivity during the evolution experiment. (****A) A histogram of all communities' ratio between their productivity at generation ~400 to generation ~10 as the ratio between the OD they reached at the end of each growth cycle. To reduce noise, OD trajectories were smoothed by moving means with a window of three, and the OD of each unique assembly was averaged across all replicates. (B) the log increase in OD of the different species when grown in monoculture.*

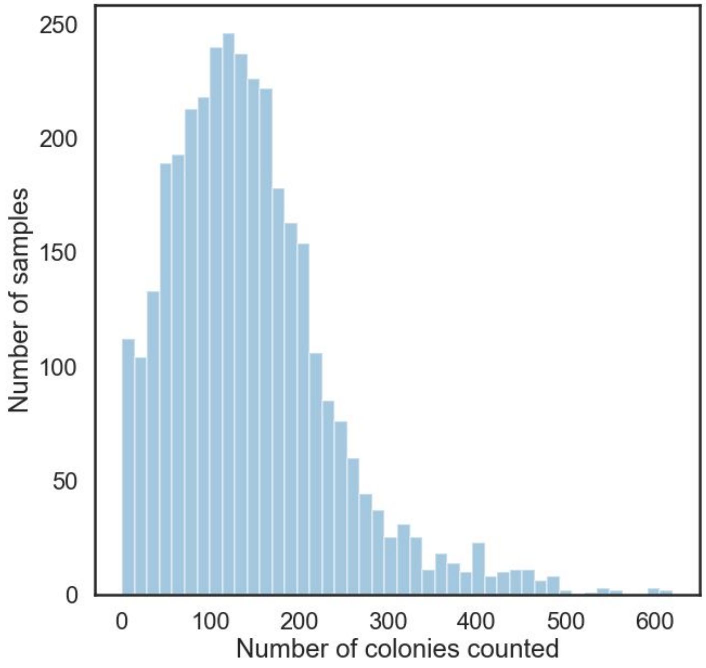


***Figure S12: The distribution of the number of colonies counted to assay community composition throughout the experiment.***

| Short Name | Species | source |
|---|---|---|
| Ea | *Enterobacter aerogenes ATCC 13048* | ATCC |
| Pa | *Pseudomonas aurantiaca ATCC 33663* | ATCC |
| Pci | *Pseudomonas citronellolis ATCC 13674* | ATCC |
| Pv | *Pseudomonas veronii ATCC 700474* | ATCC |
| Pch | *Pseudomonas chlororaphis ATCC 9446* | ATCC |
| Pf | *Pseudomans Fluroescens ATCC 506* | ATCC |
| Sm | *Serratia marcescens ATCC 13880* | ATCC |
| Ab | *Acinetobacter baylyi ATCC 3330* | ATCC |
| Fj | *Flavobacterium johnsonia strain UW101* | ATCC variant[39] |
| H77 | *Arthrobacter phenanthrenivorans* | Tomato pot |
| H79 | *Delftia lacustris* | Tomato pot |
| H82 | *Pseudomonas putida* | Tomato pot |
| H97 | *Pseudomonas pseudoalcaligenes* | Tomato pot |
| IN63 | *Rhodococcus soli* | Wheat plot |
| IN65 | *Pseudomonas alcaligenes* | Wheat plot |
| IN72 | *Pseudomonas sp. BSP5* | Wheat plot |

***Table S1: strains used in this study**. Environmental isolates are assigned to taxonomy by 16S sequence.*

| Sample | Replicates | Sample | Replicates | Sample | Replicates | Sample | Replicates |
|---|---|---|---|---|---|---|---|
| Ab | 6 | H77 | 4 | Pch_Pci_IN63 | 9 | Pf_Ab_IN72 | 3 |
| Ab_Fj | 6 | H79 | 4 | Pch_Pci_IN65 | 3 | Pf_Fj | 6 |
| Ab_H79 | 3 | H79_Fj | 6 | Pch_Pci_Pf | 3 | Pf_IN72 | 9 |
| Ab_IN72 | 9 | H79_IN65 | 3 | Pch_Pci_Pv | 3 | Pf_Sm | 9 |
| Ea | 6 | H79_IN72 | 3 | Pch_Pf | 9 | Pf_Sm_Ab | 9 |
| Ea_H79 | 3 | H82 | 4 | Pch_Pf_Ab | 3 | Pv | 6 |
| Ea_H82 | 6 | H82_H97 | 3 | Pch_Pf_Sm | 9 | Pv_H77 | 2 |
| Ea_H82_H97 | 3 | H97 | 4 | Pch_Pv_Ab | 3 | Pv_Pf | 6 |
| Ea_H97 | 3 | IN63 | 6 | Pch_Pv_IN65 | 3 | Pv_Pf_Ab | 3 |
| Ea_IN72 | 3 | IN65 | 4 | Pch_Pv_Pf | 3 | Pv_Pf_Sm | 3 |
| Ea_Pa | 9 | IN65_IN72 | 3 | Pch_Pv_Sm | 3 | Pv_Sm | 3 |
| Ea_Pa_IN72 | 3 | IN72 | 6 | Pch_Sm | 18 | Pv_Sm_Ab | 3 |
| Ea_Pa_Pv | 10 | Pa | 6 | Pch_Sm_Ab | 3 | Pv_Sm_IN65 | 3 |
| Ea_Pch | 9 | Pa_IN63 | 9 | Pch_Sm_H79 | 3 | Sm | 6 |
| Ea_Pch_H79 | 3 | Pa_Pci | 3 | Pch_Sm_IN65 | 3 | Sm_Ab | 3 |
| Ea_Pch_Pci | 2 | Pa_Pci_Fj | 3 | Pci | 6 | Sm_Ab_H79 | 3 |
| Ea_Pch_Pf | 7 | Pa_Pci_IN72 | 3 | Pci_Fj | 3 | Sm_H79 | 3 |
| Ea_Pch_Pv | 3 | Pa_Pci_Pv | 3 | Pci_H77 | 3 | Sm_H82 | 3 |

| | | | | | | | |
|---|---|---|---|---|---|---|---|
| Ea_Pci | 6 | Pa_Pv | 9 | Pci_IN63 | 9 | Sm_IN65 | 6 |
| Ea_Pci_IN72 | 2 | Pa_Pv_IN72 | 1 | Pci_IN65 | 3 | Sm_IN65_H79 | 3 |
| Ea_Pci_Pf | 3 | Pch | 6 | Pci_IN72 | 3 | | |
| Ea_Pci_Pv | 2 | Pch_Ab | 3 | Pci_Pf | 3 | | |
| Ea_Pf | 9 | Pch_Ab_H79 | 3 | Pci_Pf_IN72 | 3 | | |
| Ea_Pf_IN72 | 3 | Pch_H79 | 3 | Pci_Pv_IN65 | 3 | | |
| Ea_Pv | 9 | Pch_IN63 | 8 | Pci_Pv_IN72 | 3 | | |
| Ea_Pv_IN72 | 3 | Pch_IN65 | 6 | Pci_Pv_Pf | 3 | | |
| Ea_Pv_Pf | 6 | Pch_IN65_H79 | 3 | Pf | 6 | | |
| Fj | 4 | Pch_Pci | 9 | Pf_Ab | 18 | | |

***Table S2: Communities and replicates****. The different unique assemblies and the corresponding number of replicates used for each. These numbers include only replicates that were used for the final analysis, and do not include replicates that got contaminated and were excluded.*
